# Darwinian properties and their trade-offs in autocatalytic RNA reaction networks

**DOI:** 10.1101/726497

**Authors:** Sandeep Ameta, Simon Arsène, Sophie Foulon, Baptiste Saudemont, Bryce E. Clifton, Andrew D. Griffiths, Philippe Nghe

## Abstract

Discovering autocatalytic chemistries that can evolve is a major goal in systems chemistry and a critical step towards understanding the origin of life. Autocatalytic networks have been discovered in various chemistries, but we lack a general understanding of how network topology controls the Darwinian properties of variation, differential reproduction, and heredity, which are mediated by the chemical composition. Using barcoded sequencing and droplet microfluidics, we establish a landscape of thousands of networks of RNAs that catalyze their own formation from fragments, and derive relationships between network topology and chemical composition. We find that strong variations arise from catalytic innovations perturbing weakly connected networks, and that reproduction increases with global connectivity. These rules imply trade-offs between reproduction and variation, and between compositional persistence and variation along trajectories of network complexification. Overall, connectivity in reaction networks provides a lever to balance variation (to explore chemical states) with reproduction and heredity (persistence being necessary for selection to act), as required for chemical evolution.

## Main Text

Autocatalytic reaction networks, where chemicals collectively catalyze the synthesis of each other, are a crucial ingredient for the emergence of evolution, as they enable the maintenance and reproduction of out-of-equilibrium chemical states. Experimentally, autocatalytic chemistries built from a broad range of molecules including organic^1^ and inorganic molecules^2,3^, macrocycles^4^, peptides^5^, DNAs^6^, and RNAs^7,8^, have been described. Theoretically, it has been shown that autocatalytic reaction networks can emerge from random pools^9–12^ and several scenarios have been proposed for how they could have supported early modes of evolution^10,13–17^. Although the proposed dynamics and suggested chemical embodiments differ, these scenarios have in common the possibility to form a diversity of autocatalytic systems, with evolution arising from transitions between such systems. Consistently, recent experiments indicate that RNA replicases (RNA-dependent RNA-polymerase ribozymes) may have emerged as components of such networks^18^. This is significant as the spontaneous appearance of an RNA replicase with sufficient processivity to allow self-replication and enough fidelity to avoid an error catastrophe^19,20^ seems unlikely, given the length (>165 nt) and structurally complexity of known replicases^8,21–23^.

Nevertheless, sustaining evolution in reaction networks is by no means trivial as networks which could support the Darwinian properties of variation, differential reproduction, and heredity are not known. Indeed, these properties are mediated by chemical compositions (the chemical species present and their concentrations), rather than by the copying of a sequence, as in the template-based replication process observed in contemporary biology. Furthermore, evolution in reaction networks may be constrained by trade-offs between these properties. For instance, robustness to environmental perturbations and persistence of compositions are necessary for selection to act, but must be balanced with variation to explore novel states.

Here, by developing a method to generate a wide diversity of reaction networks and measure their compositional trajectories, we report the first large scale study of the relationship between reaction network topology and Darwinian properties in a prebiotically relevant experimental model. Analysis of this landscape reveals that variation and reproduction are controlled by a few parameters characterizing network connectivity and perturbations in terms of specific interactions. In turn, these molecular rules are found to impose trade-offs between Darwinian properties across trajectories of network growth by accretion of novel species.

## Results and discussion

We studied a model of autocatalytic RNAs derived from the group I intron^24^ of the *Azoarcus* bacterium, where fragments, denoted WXY and Z, assemble into non-covalent complexes that catalyze the formation of more efficient covalent ribozymes^7,25^ (denoted WXYZ, Fig. 1a), which in turn catalyze, with higher efficiency, the formation of further covalent ribozymes^25^. Three nucleotide long sequences, called internal guide (IGS) and target (tag), located at extremities of the WXY fragments^25^ (Fig. 1a, top left), determine catalytic specificity by base-pairing between a ribozyme IGS and a fragment tag, combinations of which yield networks of diverse connectivity (Fig. 1a, bottom center)^26^. In a coarse-grained representation of these networks by directed graphs (Fig. 1a, top right), a node represents both non-covalent and covalent ribozyme species with the same IGS and tag, and a directed edge points from an upstream ribozyme species to the downstream ribozyme species whose formation it catalyzes.

**Fig. 1.**
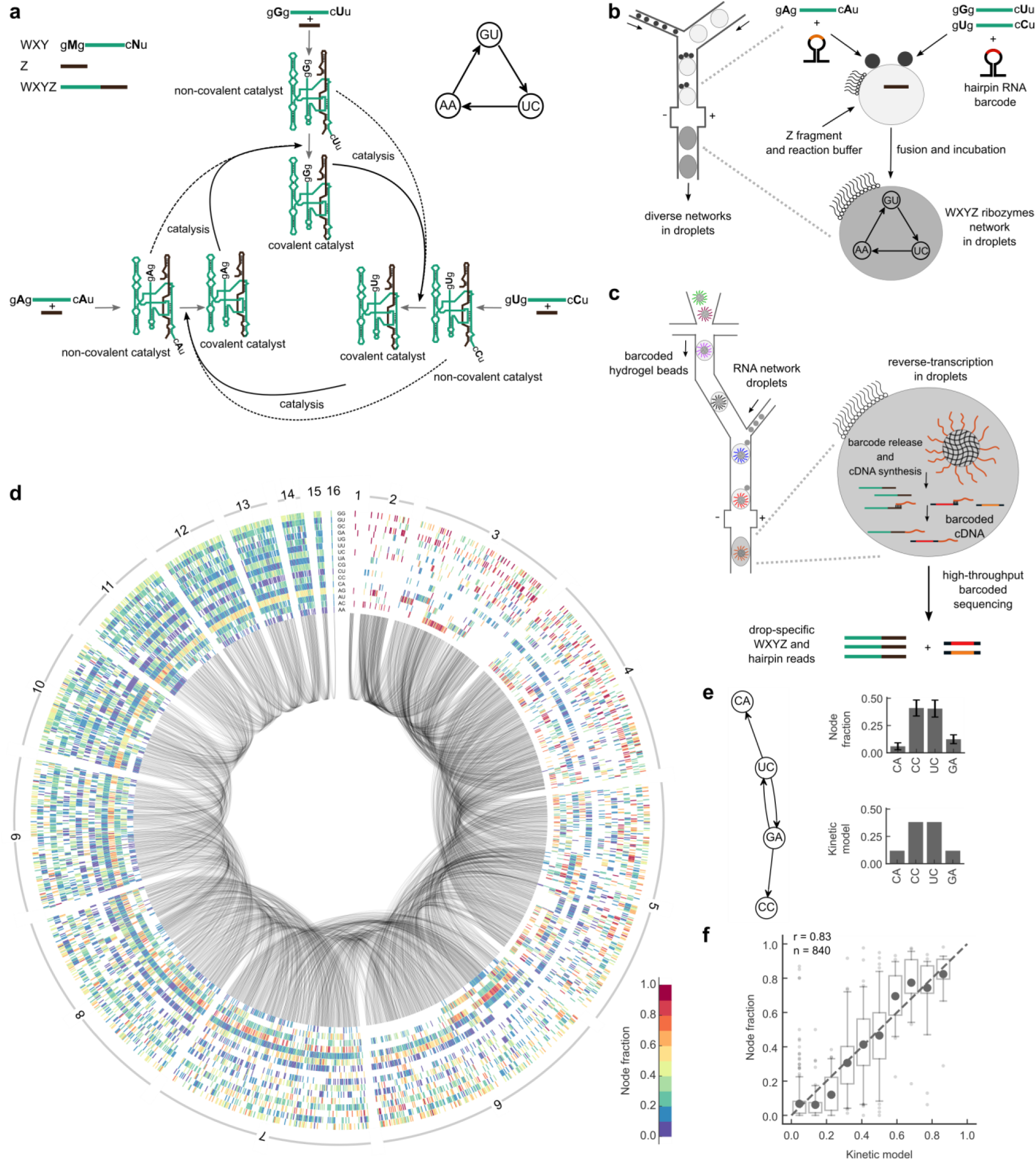
Experimental set-up and RNA network compositional landscape. **a**, Top left: RNA fragments, named WYX and Z, can be assembled into full length ribozymes WXYZ (both non-covalent and covalent catalysts). WYX fragments each comprise 3 nucleotide long IGS and tag sequences, respectively denoted gMg and cNu, where M, N can be varied, resulting in 16 different combinations^26^. Bottom center: The base-pairing interactions between M and N determine the specificity of full-length catalysts for their substrate fragments. In this example, each of the 3 different MN ribozyme species catalyzes the formation of another species from fragments provided as substrates, forming an autocatalytic cycle. Dotted and bold arrows show catalysis carried out by non-covalent and covalent ribozymes, respectively. Note that, for clarity, only Watson-Crick IGS-tag interactions are shown here. Any other non-Watson pair will have a weaker but non-zero activity (Supplementary Fig. 6C), therefore all networks possess some degree of autocatalysis. Top right: a simplified network representation shows catalytic species as nodes with their MN identity, and catalytic interactions as directed edges (corresponding to the dotted and bold black arrows of the bottom center panel). **b**, Random sets of ~3 droplets (small black disks), each containing different mixtures of WXY fragments, are fused by electrocoalescence in a microfluidic device with a droplet containing Z fragments and the reaction buffer (larger gray disks). This combinatorial process leads to a large diversity of catalytic networks, each contained in a different droplet (Supplementary Methods). Substrate fragments are initially co-diluted with non-reactive hairpin RNA reporters that later allow the initial composition to be retrieved by sequencing. **c**, Droplet-level barcoded sequencing: each RNA-containing droplet is analyzed after incubation by fusing a sample of it to another droplet containing a hydrogel bead carrying DNA barcodes. These barcodes, specific to each bead, serve as primers for the reverse transcription of the RNAs, including the ribozymes formed during incubation, and the hairpin RNA reporters to identify initial condition. **d**, Compositional landscape of the 1837 unique measured networks^36^: each ray is made of 16 boxes corresponding to the 16 possible MN species. A blank box means that the MN ribozyme is absent in this network, as determined from hairpin sequencing. Otherwise the box color indicates the measured fraction of the MN species. Grey arcs connect networks that differ from each other by a single node (addition or removal). **e**, Left: schematic of an *Azoarcus* network. Right: top, experimentally determined node fractions for the same network (error bars are ±1 s.d. over n=425 droplets); bottom, theoretical results with the kinetic model. **f**, Fraction obtained with the kinetic model versus measured fraction for every node that is a member of a 4 node-network. Bins with less than 10 points are discarded. Dark gray dots are mean values, quartile boxplots have 5th - 95th percentiles whiskers with flier points. The dotted grey line is the identity line. P-value < 0.001.

We developed a method to generate and measure a high diversity of such RNA reaction networks, using droplet microfluidics coupled to barcoded Next-Generation Sequencing (Fig. 1b, c and Supplementary Fig. 1). It consists of first producing a library of 5 pL droplets containing 24 different combinations of the 16 WXY fragments (Supplementary Table 1), together with hairpin RNA reporters that do not react, but enable later identification of the mixture of WXY fragments by sequencing. The 24 initial combinations were selected among a random sample by minimizing their mutual overlap while covering as evenly as possible network sizes from 1 to 16 species after combinatorial droplet fusion (Supplementary Methods). Random sets comprising 1 to 5 droplets of this initial library were then electrocoalesced^27^ with a 50 pL droplet containing the reaction buffer and Z fragments (Fig. 1b). After incubation at 48°C for 1 hour, the composition of each droplet was analyzed by droplet-level RNA sequencing adapted from single-cell transcriptomics^28^. Hydrogel beads carrying cDNA primers with a barcode specific to each bead (Supplementary Fig. 2) were encapsulated one-by-one in droplets, which were then fused with single RNA containing droplets, the primers released and cDNA synthesis performed in the droplet (Fig. 1c, Supplementary Fig. 1, and Supplementary Methods). Barcoded cDNAs were recovered and sequenced. Reads from the same droplet carry the same barcode (Supplementary Fig. 3a, b).

The initial combinations of fragments (encoded by hairpin RNA reporters, Supplementary Fig. 3, 4), and the final fraction of ribozymes produced during incubation were determined in 20,038 droplets, comprising 1,837 unique networks, with on average 11 replicates each (Fig. 1d and Supplementary Fig. 5a), indicating a mean precision of ~6% in species fractions (Supplementary Fig. 5b). Repeatability was tested with a full experimental replicate (r=0.84 between species fractions; Supplementary Fig. 5c). Furthermore, controls with mixtures of droplets containing defined sets of ribozymes in known proportions showed that 87% of sequenced droplets contained a single set or ribozymes and a high correlation (r = 0.91) between measured and expected ribozyme concentrations (Supplementary Fig. 5d, e).

We observed heterogeneous patterns of covalent ribozyme accumulation, demonstrating the existence of interdependences between reaction network components (Fig. 1d). Indeed, if every ribozyme species were reacting independently, the relative rank between any pair of ribozyme species would be conserved across all networks. On the contrary, we observed that for 63% of ribozyme pairs, their relative ranking differed in at least 10% of the networks (Supplementary Fig. 6a), extending previous findings of non-conserved ranking in a small set of such networks^29^. This diversity in species fractions in networks was well predicted by a kinetic model without fitting parameters (Supplementary Information) by adding the first order effective catalytic rates of non-covalent and covalent ribozymes measured independently for individual IGS-tag pairs (r=0.83, p<10^−3^, Fig. 1e, f and Supplementary Fig. 6b-d).

Comparing networks generated from distinct substrate sets also revealed differences in growth, quantified as the concentration of covalent ribozyme accumulated during the reaction relative to hairpin reporters of known concentration (Fig. 2a): yield varied 10-fold across networks comprising between 2 and 15 species (see Supplementary Table 2), and up to 6-fold within networks of the same size but with different topologies (Fig. 2a).

**Fig. 2.**
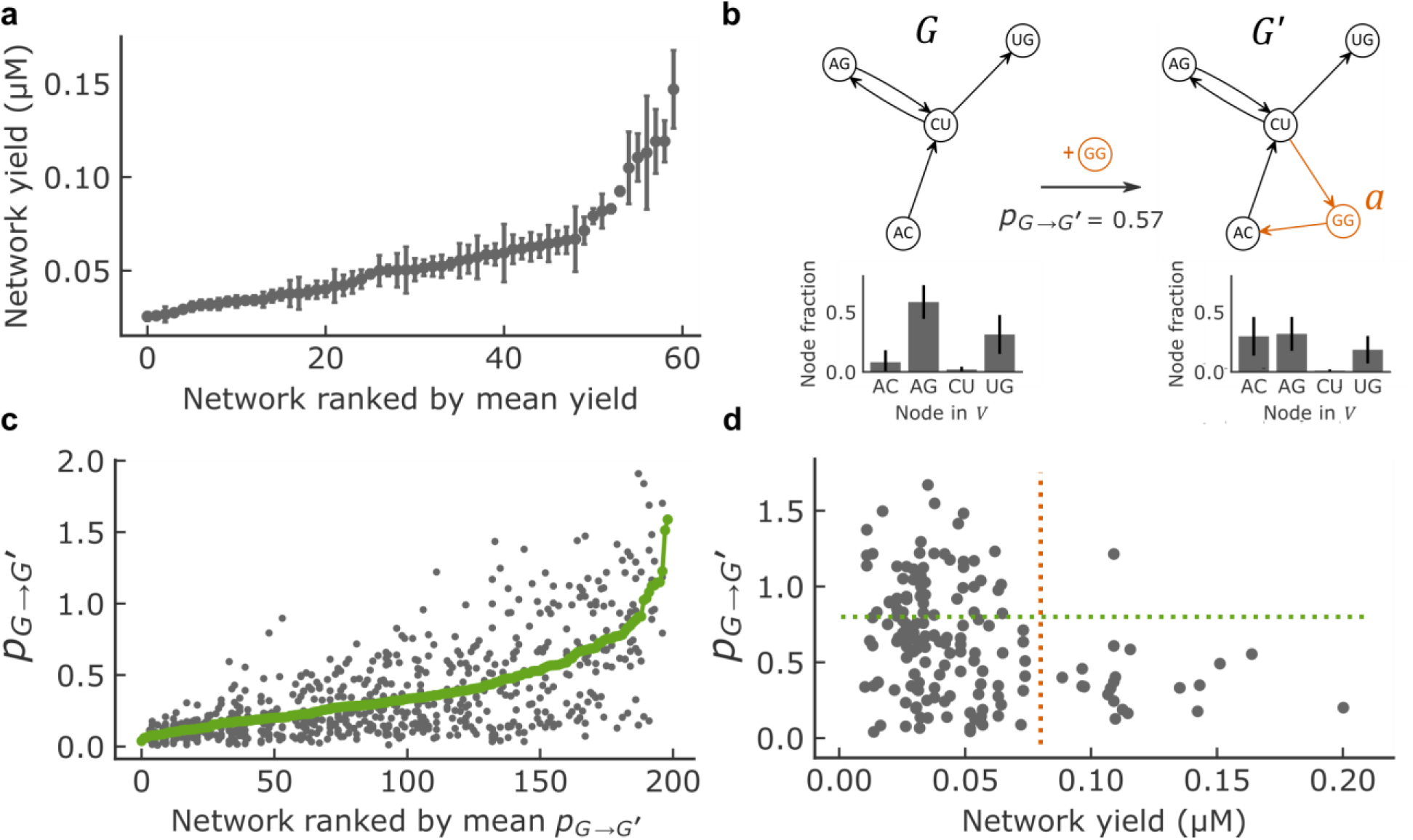
Network growth and perturbation. **a**, Network total yield (concentration of WXYZ catalysts in µM) distribution of networks with four nodes and at least 10 replicates (for data on all network sizes see Supplementary Table 2). Networks are ranked (x-axis) based on the mean across network replicates of the total WXYZ concentration (y-axis). The error bars show the Standard Error of the Mean. See Methods for further details on network yield determination. **b**, Example showing the addition of node *a* (GG, orange) to network *G* resulting in network *G’*. Top: schematic. Bottom: experimentally determined node fractions for the same network (error bars are ±1 s.d. over n=109 droplets (*G*) and n=15 droplets (*G’*). **c**, Perturbation distribution of networks with four nodes. Networks are ranked (x-axis) based on the mean perturbation (y-axis). Each position on the x-axis corresponds to a certain network *G*, for which the perturbation has been computed for all possible single node additions present in the dataset, leading to several y-axis values corresponding to different *G’* networks. The green line is the average perturbation taken over all *G*’ networks for each network *G*. **d**, Perturbation *p*_*G→G′*_ for networks with 3, 4 and 5 nodes plotted against network yield in µM for perturbations involving the addition of a novel catalysts (with G/C as IGS) with at least one target. The number of strongly perturbable (*p*_*G→G′*_ > 0.8, above the green dotted line) and high yield networks (> 0.08 µM, right hand side of red dotted line) is very low compared to the null hypothesis of independence of perturbability and yield (one-sided Fisher exact test, N=162, odds ratio < 0.1, 95% confidence interval 0.01-0.66, p-value=1.6.10^−3^).

To quantify variations in response to the addition of fragments allowing the formation of a novel ribozyme, we compared networks *G* and *G’* differing by a single node *a* (networks connected by a curved line in Fig. 1d). Denoting *V* the set of nodes common to *G* and *G’*, the total perturbation is defined as *p*_*G→G′*_ = ∑_*ν∈V*_ |*y′*_*ν*_ − *y*_*ν*_| where *y*_*ν*_ and *y′*_*ν*_ are the respective fractions of species ν in *G* and *G’*, both normalized within *V* (Fig. 2b). By definition, the maximum is *p*_*G→G′*_ = 2 and is reached for a full switch in species composition, for example going from (*y*_*u*_, *y*_*ν*_) = (0, 1) to (*y′*_*u*_, *y′*_*ν*_) = (1, 0) in *V* = {*u*,*ν*}. Averaging for each network the effect of all perturbations (green line, Fig. 2c) revealed large differences from marginally (mean 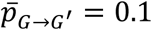) to strongly perturbable networks 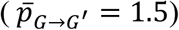. Furthermore, we observed a trade-off between growth and variation (Fig. 2d), implying that high yield networks tend to be poorly perturbable.

We next aimed to interpret the diversity of observed responses based on network connectivity. The kinetic model does not provide such a direct interpretation as it consists of a mixture between two regimes, where either only non-covalent or only covalent ribozymes are active. For non-covalent ribozymes only, species fractions should be predicted by in-degree centrality, a network-theoretic measure which accounts only for catalysis by directly upstream ribozymes^30^. In contrast, for covalent ribozymes only, species fractions should be predicted by eigenvector centrality, which accounts for longer catalytic chains^30^. Despite these differences, the in-degree centrality was found to be a good approximation of the eigenvector centrality for our dataset (R=0.83, p-value<10^−5^, Supplementary Fig. 7a), as is already known in general^31^, except for ribozymes strongly catalyzing their own production, whose fraction is underestimated (13% of the dataset, Supplementary Fig. 7b).

The in-degree centrality approximation allowed an analytical derivation of the perturbations (supplementary information),

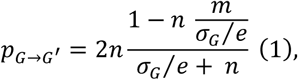

with 4 control parameters (Fig. 3a): *n* is the *perturbation breadth*, the number of targets of the new node *a* introduced in the network; *m* is the *catalytic novelty*, the number of catalysts already present in *G* with the same IGS as *a* (novelty being higher for lower *m*); *σ*_*G*_ is the *background strength*, the sum of all edge weights in the network, and; *e* is the *catalytic strength* of edges outgoing from *a* as determined by its IGS.

**Fig. 3.**
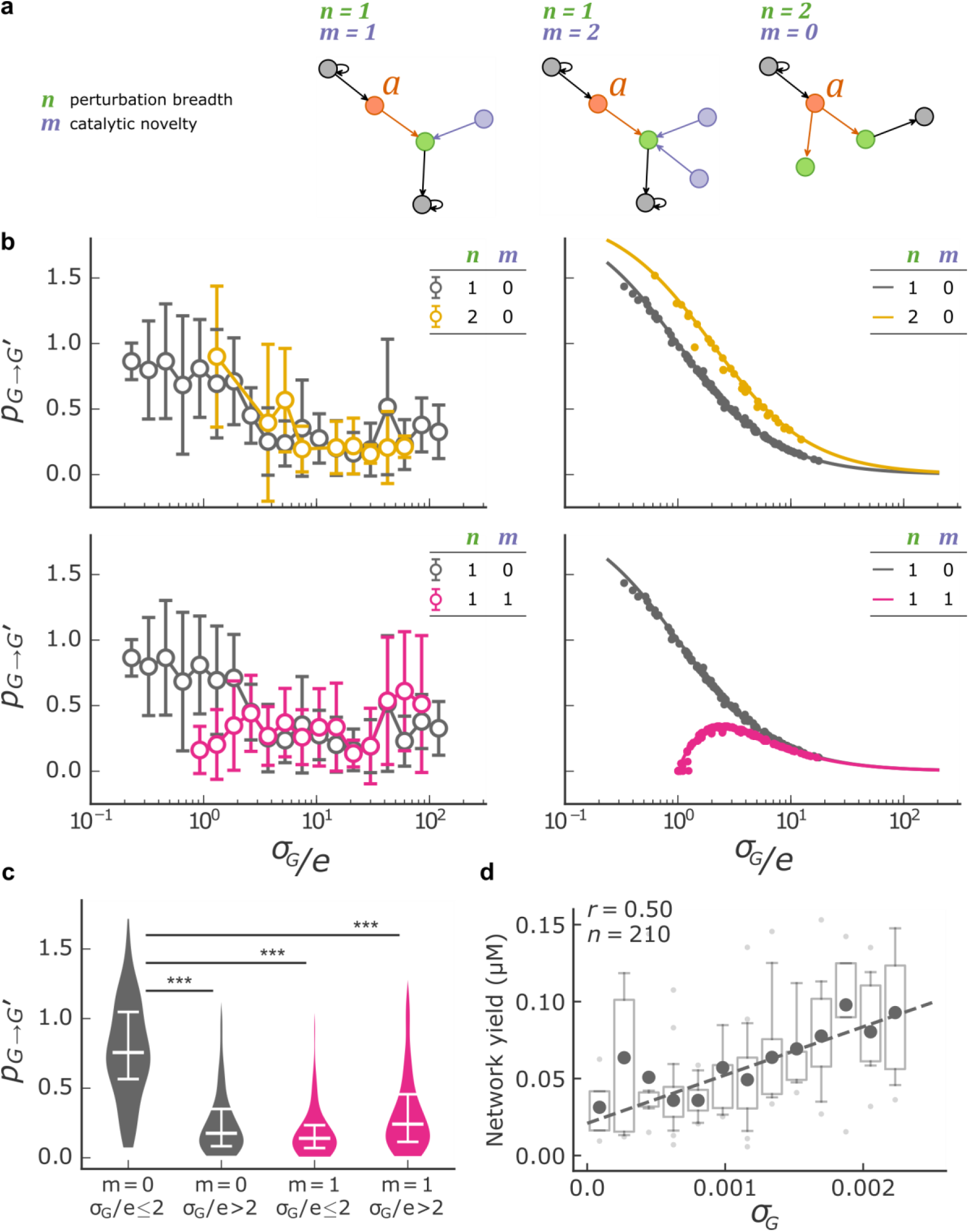
Network perturbation analysis. **a**, Schematic of three situations where the parameters *n*, the number of targets of added node *a*, and *m*, the number of nodes sharing the same IGS as *a*, have different values. **b**, The perturbation *p*_*G→G′*_ is plotted against *σ*_*G*_/*e* for different values of the parameters *m* and *n* for both the experimental data (left, mean and standard deviation per bin, bins with less than 3 points are discarded) and results predicted by in-degree centrality using the analytical expression (right, solid line) and with fractions predicted by eigenvector centrality (right, solid circles) for the networks with between 3 and 6 nodes before addition. Results for other parameter values are shown in fig. S8. **c**, Violin plot (with the median and interquartile range in white) of *p*_*G→G′*_ with catalytic innovations (*m* = 0, grey) or without catalytic innovations (*m* = 1, pink) at high and low values of *σ*_*G*_/*e*. Mann–Whitney U test p-value is reported (***: p-value<0.001). **d**, Network yield (µM, see Methods) is plotted against *σ*_*G*_ for network with four nodes.

The influence of parameters *σ*_*G*_,*e*,*m*, and *n* on *p*_*G→G′*_ predicted by the in-degree approximation (Equation 1, continuous curves in Fig. 3b) was highly similar to those predicted by eigenvector centrality (Fig. 3b, dots), and, more importantly, were well verified experimentally (Fig. 3b, compare left and right panels). First, perturbation strength was impacted much more strongly by catalytic novelty *m* (Fig. 3b, bottom) than by perturbation breadth *n* for the values tested (*n* = 1 or 2, Fig. 3b, top). In particular, for *m* = 1, networks are robust to perturbations at both low and high connectivity *σ*_*G*_/*e* (pink, Fig. 3b). Second, Equation (1) poses the condition *σ*_*G*_/*e* ≤ *n* for significant variations, high *p*_*G→G′*_ values being observed only when *σ*_*G*_/*e* ≤ 2 (given that *n* ≤ 2, Fig. 3b, top). Overall, the highest perturbations required a catalytic innovation (*m* = 0) to be combined with a low normalized background strength (*σ*_*G*_/*e* ≤ 2) (Fig. 3c). The predictions of Equation 1 were also verified for other values of the parameters (Supplementary Fig. 8) and at the level of single nodes (Supplementary Fig. 9). In addition, the attenuation of perturbations with large *σ*_*G*_/*e* combined with the weak but significant correlation between growth and *σ*_*G*_ (R=0.5, p<10^−5^, Fig. 3d) explains part of the trade-off between growth and perturbation reported in Fig. 2d.

To test the interplay between robustness and variation in scenarios where novel species would appear either spontaneously^15^ or due to changes in substrates provided from the prebiotic milieu^32,33^, we analyzed cumulative perturbations across trajectories of network growth, starting from networks with three nodes, randomly adding one node at a time (Fig. 4a). We have seen that strong perturbations require catalytic innovations (*m* = 0), Equation 1 then reducing to *p*_*G→G′*_ = 2/(1 + *σ*_*G*_/*ne*). The ratio *σ*_*G*_/*ne* expresses a second trade-off, between the robustness induced by the background strength of the network *σ*_*G*_, and the variation induced by the novel catalyst as characterized by *n* and *e*.

**Fig. 4.**
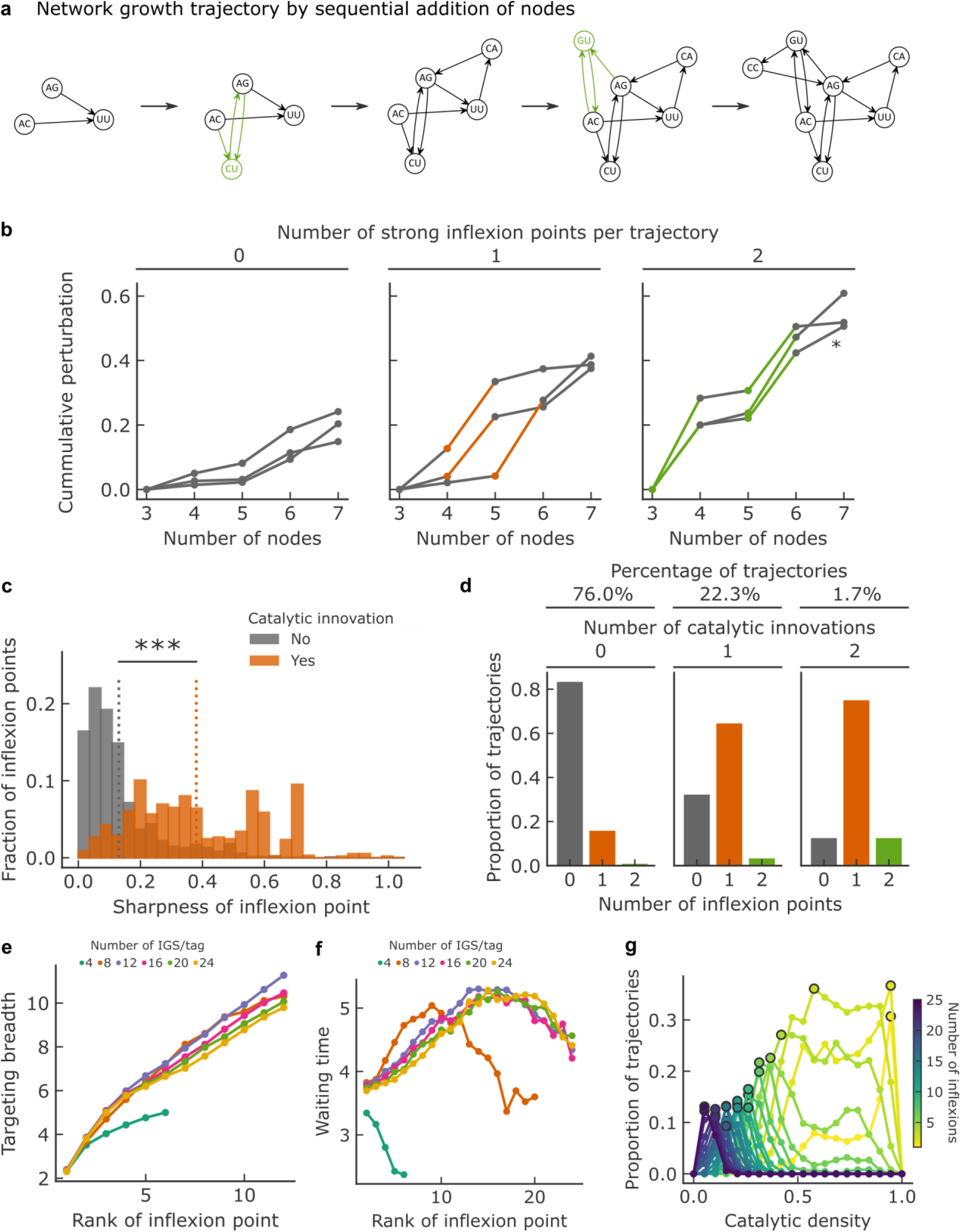
Perturbation dynamics across network growth trajectories. **a**, Example of a network growth trajectory where at each step, a new node is added. Nodes resulting in strong perturbations (see panel B) are in green, and correspond to the IGS/tag pairs C-G and G-C. **b**, Examples of measured cumulative perturbation over trajectories, plotted against the number of node additions, by number of strong inflexions points (colored). The latter are determined as the top 25% in sharpness (absolute value of the third derivative, Supplementary Fig. 10A) measured over all trajectories. The asterisk (*) denotes the perturbation trajectory of the example shown in panel **a**. **c**, Distribution of sharpness for inflexion points associated with a catalytic innovation (orange, n=4,462) or not (gray, n=934). Catalytic innovations are defined as the introduction of strong IGS/tag interactions (CG or GC, Supplementary Fig. 6B, C) that were not present until node addition. The dotted line is the mean of each distribution and the significance of the difference between the two distributions is reported (Mann–Whitney U test, p-value<0.001). **d**, Distribution of the number of inflexion points within the 75^th^ percentile sharpness per trajectory, depending on the number catalytic innovations per trajectory. **e-g**, Computational study of network growth trajectories for chemistries with varying number of IGS/tag pairs and with varying degrees of catalytic density points are determined, as before, based on their sharpness (within the 75^th^ percentile) here computed along a random sample of 1000 trajectories growing from 2 to 100 nodes. Catalytic densities are varied by random removal among the pool of all possible specific IGS/tag interactions. **e**, Targeting breath (number of targets) of catalysts causing strong perturbations, as the function of the inflexion rank. **f**, Waiting time (number of species additions) between two strong inflexions, as a function of the inflexion rank (e.g.: rank 5 is the fifth inflexion observed along a growth trajectory) **g**, Proportions of trajectories with a given number of strong inflexion points plotted against catalytic density for a chemistry comprising up to 24 different IGS/tag pairs.

As for a large perturbation to occur, *ne* must be comparable to the *σ*_*G*_ of the perturbed network, *σ*_*G*_ would roughly double after a strong perturbation, enhancing robustness to further variation. Consequently, strong variations should be followed by small ones, and result in inflexions (change in curvature) in cumulative perturbation trajectories. We quantified the corresponding inflexions by their sharpness (third derivative at inflexion, Supplementary Fig. 10a), and categorized them as strong when in the top 25% of sharpness (Fig. 4b). Comparing the distributions of sharpness for all inflexion points showed that they are, as predicted, significantly sharper for catalytic innovations (Fig. 4c). Consistently, the number of strong inflexions correlates with the number of catalytic innovations per trajectory (Fig. 4d).

Although introducing a node causing a strong perturbation buffers the resulting network against further perturbations, subsequent variations along trajectories are still possible, as exemplified in Fig. 4a and b. By computationally analyzing trajectories for an extended repertoire of specific interactions, we found that sustained variation requires perturbing species of increasing targeting breadth *n* (Fig. 4e) and increased waiting time between variation events (until the diversity of IGS/tag pairs saturates) (Fig. 4f). The former allows the ever larger *σ*_*G*_ values to be overcome, while the latter corresponds to the build-up of weakly connected nodes that can become targets. Additionally, the trade-off between *σ*_*G*_ and *ne* translates at the level of trajectories: sparse chemistries are prone to *few but strong variations*, whereas dense chemistries are prone to *many but weak variations* (Fig. 4g).

A first general constraint to achieve evolution in chemical networks resulted from theoretical studies, which established that the density of catalytic interactions must exceed a threshold for network self-reproduction^11^. Our experimental results show that connectivity in reaction networks also determines their potential for evolution, with network topology modulating both differential growth and variation in response to perturbations. Remarkably, a large range of these properties of Darwinian systems could be generated from combinations of only a few specific interactions, raising the possibility to synthesize chemistries with the capacity to evolve based on reasonably sized spaces of chemical interactions.

What then, are the requirements for reaction networks to evolve? First, we found that network-level structures constrain properties that emerge at the level of chemical compositions. Specifically, network topology imposes trade-offs between growth (favored by higher connectivity) and variation (favored by lower connectivity), and between variation (favored by lower connectivity) and robustness to perturbation (favored by higher connectivity). These considerations are directly relevant to scenarios where early evolution is driven by environmental heterogeneity^32,33^. Indeed, in prebiotic scenarios, such as Dynamical Kinetic Stability^34^, evolution depends on a balance between persistence of chemical compositions, thus robustness to perturbations, and exploration of novel compositions, thus susceptibility to perturbations. Indeed, these evolutionary trade-offs and the connectivity rules imply a ‘Goldilocks’ range (neither too high not too low) in the density of catalytic interactions for evolution to be possible.

Second, the observed dynamics points to the non-obvious role of self-assembly. Self-assembly has been demonstrated to robustly drive self-reproduction in other systems, as for example in tubular assemblies of molecules^4^. In our case, catalysis from self-assembled molecules (non-covalent ribozymes) enables the onset self-reproduction, but leads to relaxation of chemical compositions toward states determined by the substrates, rather than by the covalent catalysts transmitted from pre-existing networks (Supplementary Fig. 11), thereby limiting heredity. Reducing catalysis by self-assembled complexes consequently appears to be a critical factor to obtain multiple states with sufficient heredity to evolve.

Ultimately, provided that connectivity is balanced, heredity mechanisms are strong enough, and reactions are compartmentalized^35^, autocatalytic networks may provide a route by which novel chemical states can be explored and selected in a Darwinian manner^15^, paving the way for template-based replication, as observed in living systems today.

## Acknowledgments

The authors thanks Eörs Szathmáry for providing critical feedback on the manuscript. Authors also thank Matthew Deyell for the help in designing droplet barcodes indexes, and Estelle Mendes for her technical assistance in the realization of the experiments

## Author contributions

S. Ameta, S. Arsène, A. D. Griffiths, P. Nghe conceived the idea and designed the study. S. Ameta, S. Arsène carried out all the experiments. S. Ameta developed droplet-microfluidic experimental set-up, droplet sequencing and molecular biology protocols. S. Arsène developed custom software pipeline for sequence data treatment and data analysis, developed perturbation approach and carried out numerical simulations. S. Foulon and B. Saudemont contributed to the barcoded hydrogel bead development. B. E. Clifton carried out the kinetic measurements for *Azoarcus* self-assembly. S. Ameta, S. Arsène, A. D. Griffiths, P. Nghe interpreted the results and wrote the paper

## Data and materials availability

The dataset of species fractions for all the networks is provided as Supplementary Dataset. Other data and code are available from the corresponding authors on request.

## Supplementary Materials

Materials and Methods

Supplementary Figures 1-11

Supplementary Tables 1-2

Supplementary Movies 1-2

Supplementary Dataset 1-3

Supplementary References

